# Subthalamic nucleus neurons encode syllable sequence and phonetic characteristics during speech

**DOI:** 10.1101/2023.12.11.569290

**Authors:** W.J. Lipski, A. Bush, A. Chrabaszcz, D.J. Crammond, J.A. Fiez, R.S. Turner, R.M. Richardson

**Affiliations:** Department of Neurobiology, University of Pittsburgh, Pittsburgh, PA, USA, 15260; Brain Modulation Lab, Department of Neurosurgery, Massachusetts General Hospital, Boston, MA, USA, 02114; Harvard Medical School, Boston, MA, USA 02115; Department of Psychology, University of Pittsburgh, Pittsburgh, PA, USA, 15260; Center for the Neural Basis of Cognition, Pittsburgh, PA, USA, 15260; Learning Research and Development Center, University of Pittsburgh, Pittsburgh, PA, USA, 15260; Department of Neurological Surgery, University of Pittsburgh, Pittsburgh, PA, USA, 15260

**Keywords:** speech, sequence, electrophysiology

## Abstract

Speech is a complex behavior that can be used to study unique contributions of the basal ganglia to motor control in the human brain. Computational models suggest that the basal ganglia encodes either the phonetic content or the sequence of speech elements. To explore this question, we investigated the relationship between phoneme and sequence features of a spoken syllable triplet and the firing rate of subthalamic nucleus (STN) neurons recorded during the implantation of deep brain stimulation (DBS) electrodes in individuals with Parkinson’s disease. Patients repeated aloud a random sequence of three consonant-vowel (CV) syllables in response to audio cues. Single-unit extracellular potentials were sampled from the sensorimotor STN; a total of 227 unit recordings were obtained from the left STN of 25 subjects (4 female). Of these, 113 (50%) units showed significant task-related increased firing and 53 (23%) showed decreased firing (t-test relative to inter-trial period baseline, p<0.05). Linear regression analysis revealed that both populations of STN neurons encode phoneme and sequence features of produced speech. Maximal phoneme encoding occurred at the time of phoneme production, suggesting efference copy- or sensory-related processing, rather than speech motor planning (-50ms and +175ms relative to CV transition for consonant and vowel encoding, respectively). These findings demonstrate that involvement of the basal ganglia in speaking includes separate single unit representations of speech sequencing and phoneme selection in the STN.

## 1. Introduction

Speech requires the coordination of phonetic content and precisely timed movements of the target articulators, as well as simultaneous sensory processing of the resulting vocalizations. Neural correlates of the phonetic and articulatory components of speech as well as those for coordination of sequence timing have been observed in cortical brain regions (Bohland and Guenther, 2006; Bouchard et al., 2013; Chrabaszcz et al., 2019). Recent theoretical considerations and experimental evidence, however, have also implicated the basal ganglia in both speech and sequence production (Watson and Montgomery, 2006; Herrojo Ruiz et al., 2014; Chrabaszcz et al., 2019). The basal ganglia receive cortical inputs to the striatum, where information is transferred through the direct and indirect pathways, and to the subthalamic nucleus (STN), via the hyperdirect pathway. We recently presented evidence that cortical areas responsible for speech planning, production, and perception all project directly to the STN (Jorge et al., 2022). Here, we examined whether single-unit activity of STN neurons encodes phoneme identity and syllable sequencing during speech.

Improved understanding of the contributions of the basal ganglia to speech processing has important clinical implications. Impairments in speech production are common in basal ganglia-associated disorders including Parkinson’s disease, dystonia, and stuttering (Alm, 2004; Giraud et al., 2008; Toyomura et al., 2015). At the same time, disturbances in acoustic sensory, temporal and proprioceptive processing have also been observed in patients with Parkinson’s disease (Artieda et al., 1992; Mongeon et al., 2009; Troche et al., 2012). Limited understanding of the basal ganglia’s roles in speech impedes therapeutic innovation for speech dysfunction in these patients. At the same time, investigation of the neural correlates of speech in the basal ganglia also affords an opportunity to tackle more general questions about the roles of basal ganglia in motor control and sensory processing. A large body of pre-clinical studies infer that cortico-basal ganglia circuits consolidate isolated movements into stereotyped sequence “chunks” during the execution of learned complex motor actions (Jin et al., 2014; Ruiz et al., 2014; Jin and Costa, 2015; Dhawale et al., 2021). These experiments support long-standing ideas about the roles of the basal ganglia in habit and stimulus-response learning (Miller, 1956; Graybiel, 1998), which nevertheless remain unsettled (Turner and Desmurget, 2010; Hélie et al., 2015; Wymbs and Grafton, 2015).

The GODIVA model of the neural mechanisms of speech production postulates that a cortical-basal ganglia loop circuit controls the release of planned speech sounds to the execution system so that a new motor program is initiated only upon completion of the previous program (Bohland et al., 2010). Still, no model of speech production specifically includes the STN. The oversimplified representation of the basal ganglia in current models is unavoidable, due to the relatively low temporal and spatial resolution of functional neuroimaging and the relative inaccessibility of basal ganglia nuclei for higher resolution measurements of neural activity. Previous studies from our group and others have found modulation of neural activity in the STN during speech suggesting encoding of articulatory (Chrabaszcz et al., 2019; Dastolfo-Hromack et al., 2022) or phoneme (Tankus and Fried, 2019) features, as well as signaling related to speech timing (Watson and Montgomery, 2006; Lipski et al., 2018). Here, we investigated the modulation of STN single-unit activity during the production of syllable triplets balanced for type and sequence order of phonemes. Improved understanding of how the basal ganglia modulates the uniquely human process of speech production and perception may shed light on mechanisms of basal ganglia participation in other complex human behaviors.

## 2. Methods

### Subjects

Subjects were patients with Parkinson’s disease who provided informed consent to participate in speech production tasks during DBS lead implantation surgery, after being scheduled for surgery following the recommendation of a multidisciplinary board. Twenty-five patients (4 female) were enrolled in the study, which was approved by the University of Pittsburgh Institutional Review Board (Study # 20070368). Table 1 summarizes subject characteristics.

**Table 1.** Patient Characteristics. Twenty-five subjects performed a total of 60 sessions of the syllable triplet task during simultaneous intracranial microelectrode recording from the STN. UPDRS = Unified Parkinson’s Disease Rating Scale. NR = data not in medical record.

| Subject ID | Sex | Handed-<br>ness | UPDRS<br>Speech<br>(OFF) | UPDRS<br>Total OFF<br>Score | UPDRS<br>Speech<br>(ON) | UPDRS<br>Total ON<br>Score | Age at<br>time of<br>surgery | Length of<br>diagnosis<br>prior to<br>surgery<br>(years) | Number<br>of ses-<br>sions |
| --- | --- | --- | --- | --- | --- | --- | --- | --- | --- |
| 1 | Male | Right | 0 | 25 | 0 | 10 | 69.1 | 6.5 | 4 |
| 2 | Male | Right | 1 | 29 | 1 | 16 | 59.9 | 5.6 | 3 |
| 3 | Male | Right | 1 | 17 | 0 | 3 | 51.6 | 6.8 | 3 |
| 4 | Male | Unknown | 2 | 61 | 2 | 36 | 61.8 | 10.8 | 3 |
| 5 | Male | Right | 1 | 46 | 1 | 22 | 59.6 | 4.8 | 3 |
| 6 | Male | Right | NR* | NR* | NR* | NR* | 70.8 | 5.0 | 2 |
| 7 | Male | Right | 1 | 34 | 1 | 14 | 70 | 1.1 | 3 |
| 8 | Male | Right | 1 | 30.5 | 0 | 7 | 66.1 | 8.2 | 3 |
| 9 | Male | Right | 2 | 43 | 2 | 26 | 70 | 20.3 | 4 |
| 10 | Male | Unknown | 0 | 30 | 0 | 19 | 71 | 15.4 | 3 |
| 11 | Male | Right | 1 | 42 | 1 | 24 | 75 | 6.4 | 4 |
| 12 | Male | Right | 1 | 34 | 1 | 26 | 76.8 | 0.8 | 3 |
| 13 | Female | Right | 1 | 21 | 0 | 13 | 62 | 1.9 | 4 |
| 14 | Male | Right | NR* | 40 | NR* | 18 | 71 | 6.9 | 3 |
| 15 | Male | Right | 0 | 19 | 0 | 15 | 64 | 9.3 | 3 |
| 16 | Male | Right | 1 | 38 | 1 | 20 | 67 | 5.1 | 3 |
| 17 | Male | Right | 2 | 30 | 1 | 7 | 61 | 3.7 | 2 |
| 18 | Female | Right | 1 | 37 | 1 | 27 | 63 | 5.3 | 3 |
| 19 | Male | Right | 1 | 50 | 1 | 24 | 69 | 10.6 | 2 |
| 20 | Male | Left | 0 | 33 | 0 | 30 | 57 | 3.6 | 3 |
| 21 | Female | Left | 2 | 39 | 2 | 29 | 64 | 4.1 | 4 |
| 22 | Male | Unknown | NR* | NR* | NR* | NR* | 78 | NR* | 2 |
| 23 | Female | Right | 2 | 43 | 2 | 30 | 62 | 8.7 | 2 |
| 24 | Male | Right | 2 | 39 | 2 | 26 | 70 | 4.7 | 2 |
| 25 | Male | Right | 1 | 47 | 1 | 27 | 56 | 5.8 | 3 |

### Surgery

Deep brain stimulation lead placements were performed by a single surgeon following the procedure described in (Crammond and Richardson, 2020). The DBS lead target was first defined by magnetic resonance imaging to identify the stereotaxic location of the subthalamic nucleus. Subcortical mapping using microelectrode recording was then performed to confirm and, if necessary, refine that target location. Microelectrode recordings were performed along three parallel tracks directed at the MRI-defined location of the subthalamic nucleus, using the ‘Ben Gun’ configuration (2mm spacing; central, posterior and medial tracts).

### Electrophysiological recording

Single-unit recordings were carried out using Parylene-insulated, reduced-Larsen effect tungsten microelectrodes and the Neuro-Omega recording system (Alpha Omega), as described previously (Lipski et al., 2018). Microelectrode impedances ranged from 200 to 600 kΩ. All recordings were collected from fully awake subjects more than 30 minutes after the cessation of propofol anesthesia. Figure 1A-B shows an example of an extracellular recording from the STN.

**Figure 1.**
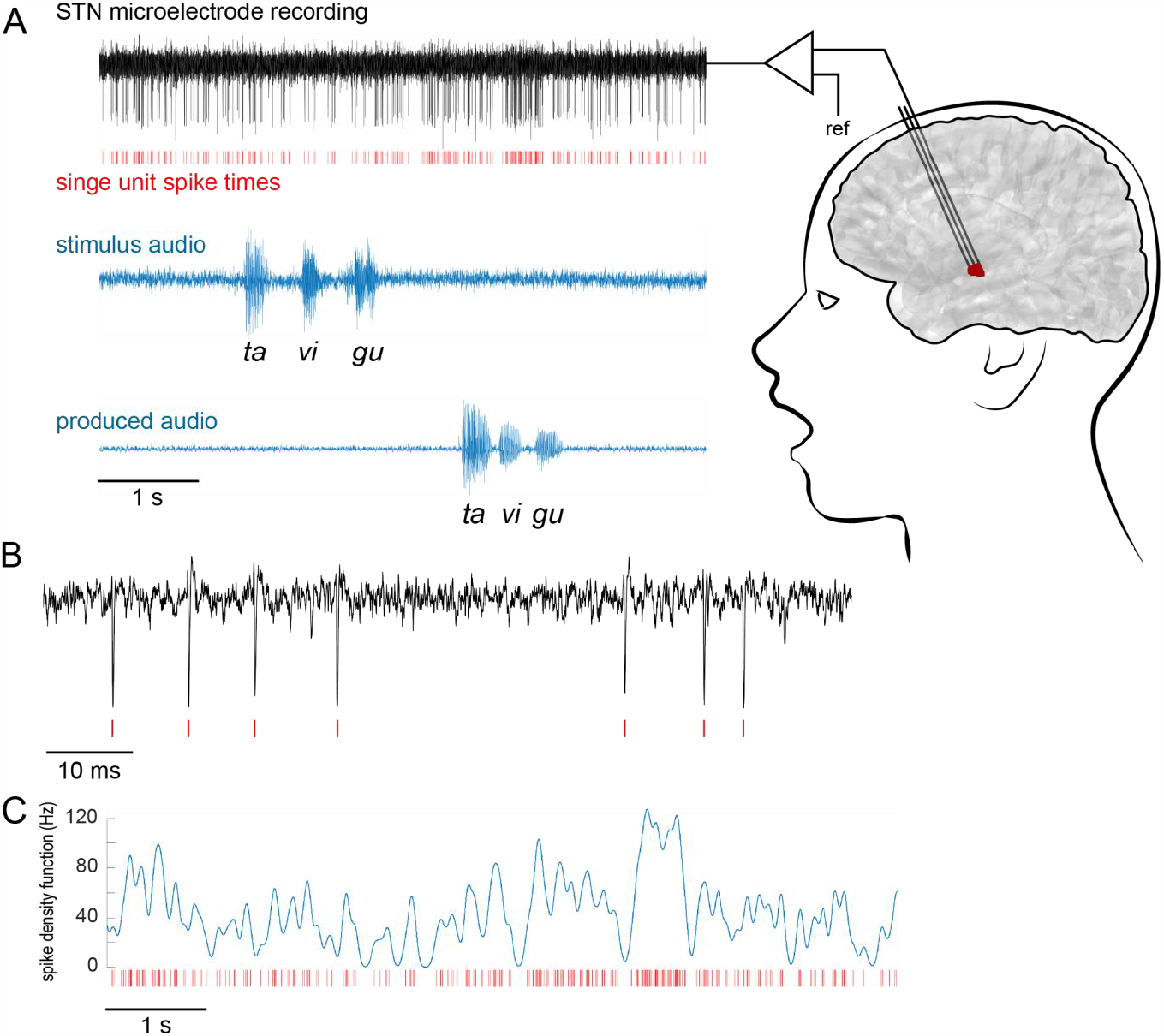
Extracellular recordings of STN neurons were performed using microelectrodes during the mapping portion of DBS implantation surgery. Patients repeated aloud syllable triplets composed of consonant-vowel pairs, which were presented as audio cues through earphones. (A) Representative recording during a single trial of the task, showing (top to bottom) the simultaneous microelectrode recording, single unit spike times, and the stimulus and produced audio waveforms. (B) Detail from (A), demonstrating action potential extracellular spike waveforms in the high-pass filtered signal. Red hash marks (realigned to spike minima) indicate sorted spikes. (C) Example of a spike density function, computed by convolving the binned single unit spike time series in (A) with a Gaussian kernel (sigma = 25 ms).

### Anatomical localization of recordings

Electrode localization was performed using the Lead-DBS toolbox (Horn and Kühn, 2015). Preoperative and postoperative magnetic resonance images were coregistered and normalized to Montreal Neurological Institute (MNI) space. MNI locations of DBS lead placements were determined from postoperative images and intraoperative microelectrode locations were calculated based on their position relative to final lead placement.

### Syllable triplet speech production task

During the awake mapping portion of the DBS electrode implantation procedure, patients were asked to listen to and then repeat auditory stimuli consisting of triplets of unique consonant-vowel syllables (Figure 1A). Each trial consisted of a one second pre-stimulus baseline period followed by the auditory presentation of a unique syllable triplet, 1.5 s in duration, via earphones (Etymotic SR4). Subjects were instructed to repeat the syllable triplet at their own pace after completion of the auditory cue. The experimenter manually initiated the next trial after completion of the subject’s vocal response plus an inter-trial interval of at least 1.0 s. Each syllable within a triplet was composed of one of four possible consonants (/v/, /s/, /t/, /g/) and one of three possible vowels (/a/, /u/, /i/). Each session consisted of up to 120 trials, in which individual syllables were presented not more than once per triplet in pseudo-randomized order and balanced with respect to the number of consonants and vowels presented at each ordinal position in the triplet sequence. Subject-produced vocal responses were recorded using an omnidirectional microphone (PreSonus, model PRM1 Precision Flat Frequency Mic, frequency response 20–20,000 Hz) and the audio signal was segmented into trials offline.

Subject responses were coded by speech pathology students trained and supervised by a speech-language pathologist using a custom-designed graphical user interface implemented in MATLAB (http://github.com/Brain-Modulation-Lab/SpeechCod-ingApp). Trials in which subjects accurately reproduced the stimulus triplet, and those in which they produced inaccurate consonant-vowel syllables (i.e. responses that did not match the stimulus triplet) but that were nevertheless composed of the target phonemes, were included in subsequent analyses.

### Spike Sorting

Microelectrode recordings were high-pass filtered at 200 Hz and a voltage threshold was applied to identify waveforms to be spike-sorted. The threshold was chosen by estimating a voltage value that includes all putative action potential spike waveforms as well as a representation of putative background and noise waveforms. Spike sorting was performed in principal components space (Offline Sorter, Plexon Inc.) (Lipski et al., 2018). Spike times were defined as the start of the waveform used for sorting. Standard criteria were used to grade the quality of unit isolation into “A-sort” and “B-sort” categories, as well as to categorize them as single-unit and multi-unit recordings, as described in (Lipski et al., 2018).

### Spike and acoustic contamination signal-to-noise estimation

To test whether acoustic contamination of the neurophysiological signals recorded in this experiment could affect our spike sorting results, we estimated the signal-to-noise ratio (SNR) of (1) sorted action potential spikes and (2) speech-related changes in voltage amplitude in the micro-electrode channels. The SNR of sorted action potential spikes was computed for each unit by taking the ratio of unit spike to unsorted waveform peak-to-peak amplitudes. The SNR of speech-related changes in voltage amplitude was estimated for each task session by first computing the root-mean-square (RMS) amplitudes of the microelectrode signal (high-pass filtered at 150 Hz) during all pre-audio stimulus baseline epochs and during all speech production epochs. The speech-related SNR was then calculated by taking the ratio of the mean RMS during speech to mean RMS in baseline epochs.

### STN unit activity during speech

Changes in neuronal firing rate around events in the syllable triplet task were estimated using two firing rate functions: (1) to test for task-related increases in activity, a millisecond-resolution spike density function (SDF) was constructed by convolving spike times with a 25 ms SD Gaussian kernel (Matlab CONV; Figure 1C), and (2) to test for task-related decreases in activity, a continuous inter-spike interval (ISI) function was constructed as a millisecond-by-millisecond representation of the current time interval between successive single-unit spikes smoothed using a 25 ms Gaussian kernel. These firing rate functions were averaged across trials around task events such as speech onset and tested for significant changes relative to activity during the one second pre-stimulus baseline period.

A unit was considered to have significantly elevated firing if the mean SDF within the test epoch exceeded a threshold level for at least 100 ms. The threshold was defined as the upper 5% of a normal distribution with a mean and σ of the baseline mean SDF across trials, Bonferroni corrected for multiple comparisons (where the number of independent observations was considered to be the duration of the epoch of interest divided by the width of the Gaussian kernel, 25 ms). Similarly, a unit was considered to have significantly reduced firing within a given epoch if the mean ISI time series exceeded a threshold ISI value for at least 100 ms. The threshold ISI value was defined as the upper 5% tail of a normal distribution with a mean and σ of the baseline mean ISI time series, Bonferroni corrected for multiple comparisons (where the number of independent observations was the mean number of ISIs within the epoch of interest) (Lipski et al., 2018).

### Regression modeling

Two separate linear regression models were fit to each unit’s neural activity during the syllable triplet task: (1) The phoneme-sequence model and (2) the phoneme model. By sequencing we understand the order of the produced syllables within a given triplet. The phoneme-sequence model estimated the mean spike density during a 200 ms window centered on the consonant-vowel transition of each syllable, using 7 categorical variables: three variables describing the 4 possible consonant phonemes (x_1_ to x_3_); 2 variables describing the 3 possible vowel phonemes (x_4_ and x_5_); and 2 variables describing the three syllable sequence positions (x_6_ and x_7_) (Figure 2A-B). The number of categorical variables within each feature category (vowel, consonant, and ordinal position in the sequence) was one less than the number of mutually exclusive conditions within that category in order to avoid over-fitting the data with redundant variables. The /vi/ consonant-vowel pair in the first position was chosen as the reference condition (x_0_), on which the rest of the feature space was built. Thus, the model can be formulated as follows:

**Figure 2.**
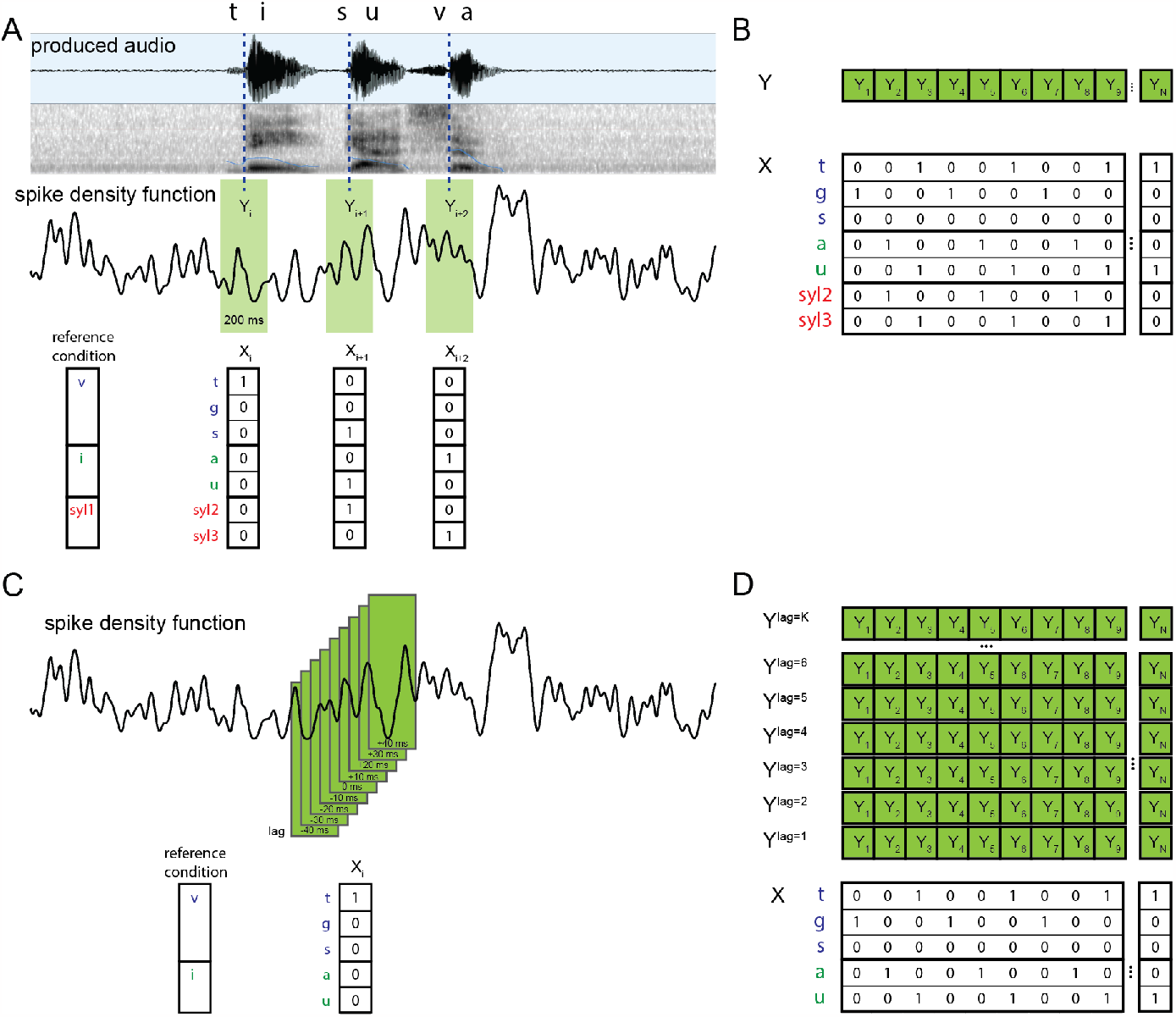
A linear regression model predicted the mean firing rate during syllable production based on phonetic and sequence features of the syllable. (A) Example of a produced audio waveform and spectrogram (top) during a single trial, and the spike density function for a simultaneously recorded STN neuron (middle). The mean firing rate within a 200 ms window centered on the consonant-vowel transition of each syllable was paired with a feature vector describing that syllable (bottom). (B) For each unit, a series of firing rates observed during all produced syllables within a task session was regressed against the corresponding syllable feature matrix. (C, D) To assess the timing of phoneme encoding relative to production, multiple regressions were performed using the mean firing rate within a series of 200 ms windows that were lagged in 25 ms increments before and after the consonant-vowel transition, in a separate analysis.

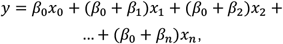

where *y* is the mean firing rate within the 200 ms window, and βn are regression coefficients in the model. The phoneme model used a reduced feature set of the 5 consonant and vowel variables (x_1_ to x_5_). This model was used to independently estimate the mean spike density during 200 ms windows at different lag times relative to the consonant-vowel transition (e.g., at lags from -1.5 s to +1.5 s in steps of 25 ms amounting to a total of 121 lag times, Figure 2C-D). Syllable sequence features were not used in the second model to avoid spurious results at large lag times due to the predictable 3-syllable task design.

R^2^ values were computed to assess the proportion of total variance in the neural activity accounted for by each model, and partial R^2^ values were evaluated for each categorical variable to assess that variable’s contribution to the model fit. The partial R^2^ for variable x_i_ was evaluated as follows:

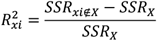

where *SSR*_*xi∉X*_ represents the sum of squared residuals from a reduced model without variable x_i_, and *SSR*_*X*_ is the sum of squared residuals from the full model. To ensure that the sum of partial R^2^ values sum to the total R^2^, and thus validate the interpretation that the partial R^2^ represents the proportion of the total explained variance, the partial R^2^ values were adjusted as follows:

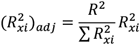

Units were included in this modeling analysis only if they maintained isolation across at least 20 trials of the speech task.

### Timing of consonant encoding

To investigate the timing of consonant encoding relative to speech onset, we examined a subset of units with a significant phoneme model fit and significant consonant encoding (N=33). Of these, we selected units with at least 8 trials containing each consonant in the first syllable position (N=24). We examined the firing rate of these units around the acoustic onset of the first syllable. Focusing on changes in the spike density function (25ms gaussian kernel) allowed us to track consonant-selective activity with a higher temporal resolution compared with the regression model (200ms moving window) and concentrating on the first syllable of the triplet eliminated the confound of activity related to preceding syllables. We examined the mean spike density functions during the production of the first syllable for consonant encoding units to test whether changes in firing rate associated with consonant encoding occurred at a consistent time relative to speech onset. Spike density functions were aligned to the acoustic onset of the first spoken consonant.

### Permutation Testing

To estimate the probability of obtaining a given R^2^ value by chance, results from each model were compared against those from 1000 surrogate models obtained by randomly permuting the feature matrix (features x trial) across trials. Thus, within the surrogate data set, the model fit was evaluated on the same spike density function but using a scrambled feature matrix. The R^2^ value from the experimental data was then compared against the distribution of surrogate R^2^ values to determine a significant model fit (p < 0.05). Similarly, the significance of the contribution of each categorical variable to the overall fit was determined by comparing the experimental partial R^2^ values against those obtained from surrogate data.

To estimate the probability of obtaining a given distribution of encoding properties among units by chance, the experimental result was compared with surrogate data obtained by randomly permuting partial R^2^ values for each categorical variable among units 100,000 times, with the permutation for each variable performed independently. The surrogate data set thus contained the same set of partial R^2^ values, but the contribution of each feature to the model was independent of other features.

### Experimental Design and Statistical Analysis

To test whether neurons in the STN encode phoneme and sequence speech features, we used a within-subject design, with 25 subjects (4 female) completing between one to four sessions of the syllable triplet production task (see Methods: Syllable triplet speech production task). We used two separate linear regression models (1) to test whether STN neural activity was modulated according to phoneme and sequence features of speech and (2) to determine the timing of neural phoneme encoding relative to acoustic speech production (see Methods: Regression modeling). Permutation testing was performed to estimate the statistical significance of model fits (see Methods: Permutation Testing). Significance level α was set at 0.05 unless otherwise noted. All analyses were carried out in MATLAB software (MathWorks Inc., Natick, Massachusetts).

## 3. Results

### Syllable triplet task performance

Subjects performed the speech task with high accuracy. Twenty-five subjects performed a total of 60 sessions of the syllable triplet task during simultaneous intracranial microelectrode recording from the STN (median sessions per subject: 2, interquartile range: 1). Subjects performed a mean of 67 trials per session that were included in the analysis. Out of a total of 6154 trials performed during STN recordings included in the analysis, 520 (8.4%) were rejected due to errors in a subject’s speech performance as judged by a speech-language pathologist.

### STN neuronal firing is modulated during triplet repetition

Consistent with previous observations (Lipski et al., 2018), many STN single units showed firing rate changes leading up to and during the production of speech. Of the 219 units recorded from 10 trials or more of the syllable triplet task, 110 (50%) met the criteria for a significant increase in firing rate (increase-type responses) around the time of onset of speech production (response window defined from the end of the auditorily presented syllable triplet cue to 500 ms after the mean triplet production offset and averaged across trials irrespective of the syllables produced). In contrast, 65 units (30%) met the criteria for a speech-related decrease from baseline (decrease-type responses). Of these, 25 units (11%) exhibited both an increase and decrease within the response window. The remaining 69 units (31%) did not show a significant firing rate change from baseline. Figure 3A-B shows example raster plots of increase- and decrease-type neurons, respectively; Figure 3C-D summarizes the timing of significant changes for all speech-modulated neurons.

**Figure 3.**
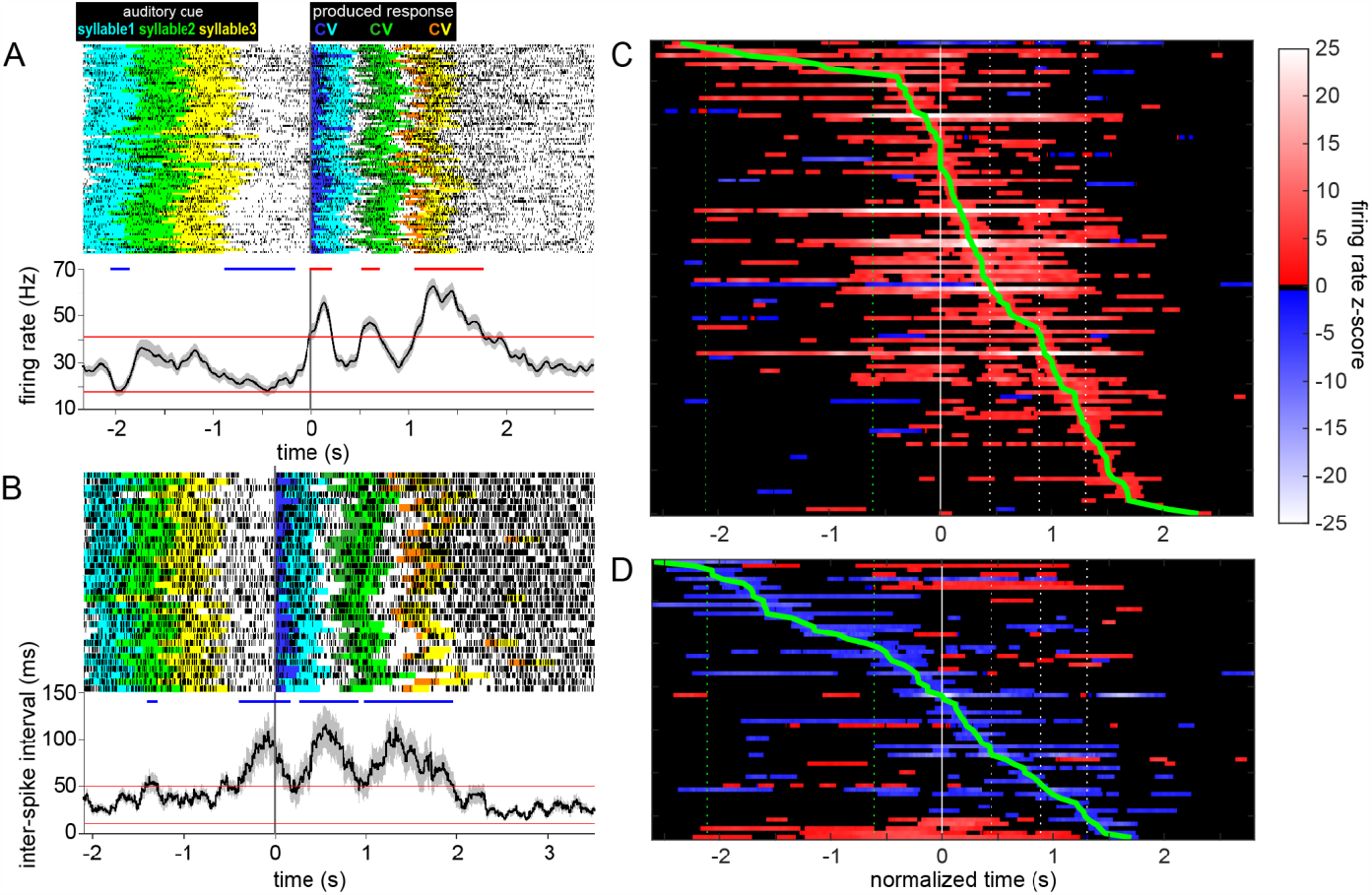
STN neuronal firing was dynamically altered during speech production. Examples of increase-type (A) and decrease-type (B) responses showing raster plots (top) and mean firing rate or inter-spike intervals (bottom). Highlighted epochs in the raster plots indicate timing of syllables in the auditory cues and each phoneme in the produced responses, as indicated above panel A. Epochs of significantly increased and decreased firing relative to baseline are indicated with red and blue horizontal bars above the histogram, respectively. Firing rate changes relative to baseline for all units showing significant increases (C) and decreases (D) within a time segment surrounding production onset. Twenty-five units had epochs of both increased and decreased firing within this segment. The time scale is normalized across units from 0.5 s before the mean cue onset until 0.5 s after the mean end of speech, aligned to production onset (t=0). The green line traces the time of maximal change in firing rate relative to baseline for each unit. Green dotted lines demarcate the timing of the auditory cue; white dotted lines indicate the onset of each produced syllable.

### STN neurons encode phoneme and sequence features of speech

The 187 units recorded over at least 20 trials of the syllable triplet task were used for the regression modelling analysis. Of these, the phoneme-sequence model resulted in a significant fit in 68 units (36%; speech-encoding units). Figure 4A-C shows mean spike density syllable raster plots of example single-units with significant model fits, demonstrating differential firing patterns across consonant, vowel and sequence features of produced speech (Figure 4A, 4B, and 4C, respectively). Spike density plots in Figures 4A and 4B show increased firing within a 200 ms window centered on the consonant-vowel transition for syllables containing the /v/ and /u/ phonemes, respectively. Figure 4C illustrates a greater amplitude and earlier onset of the firing increase during the first relative to the second and third syllable. Figure 4D-F shows corresponding regression coefficients and partial R^2^ values for each categorical variable in the model. The distribution of R^2^ values resulting from application of the phoneme-sequence regression model to all units is shown in Figure 5A, along with their anatomical distribution in the motor STN. We found no evidence for an anatomic segregation of encoding-type units.

**Figure 4.**
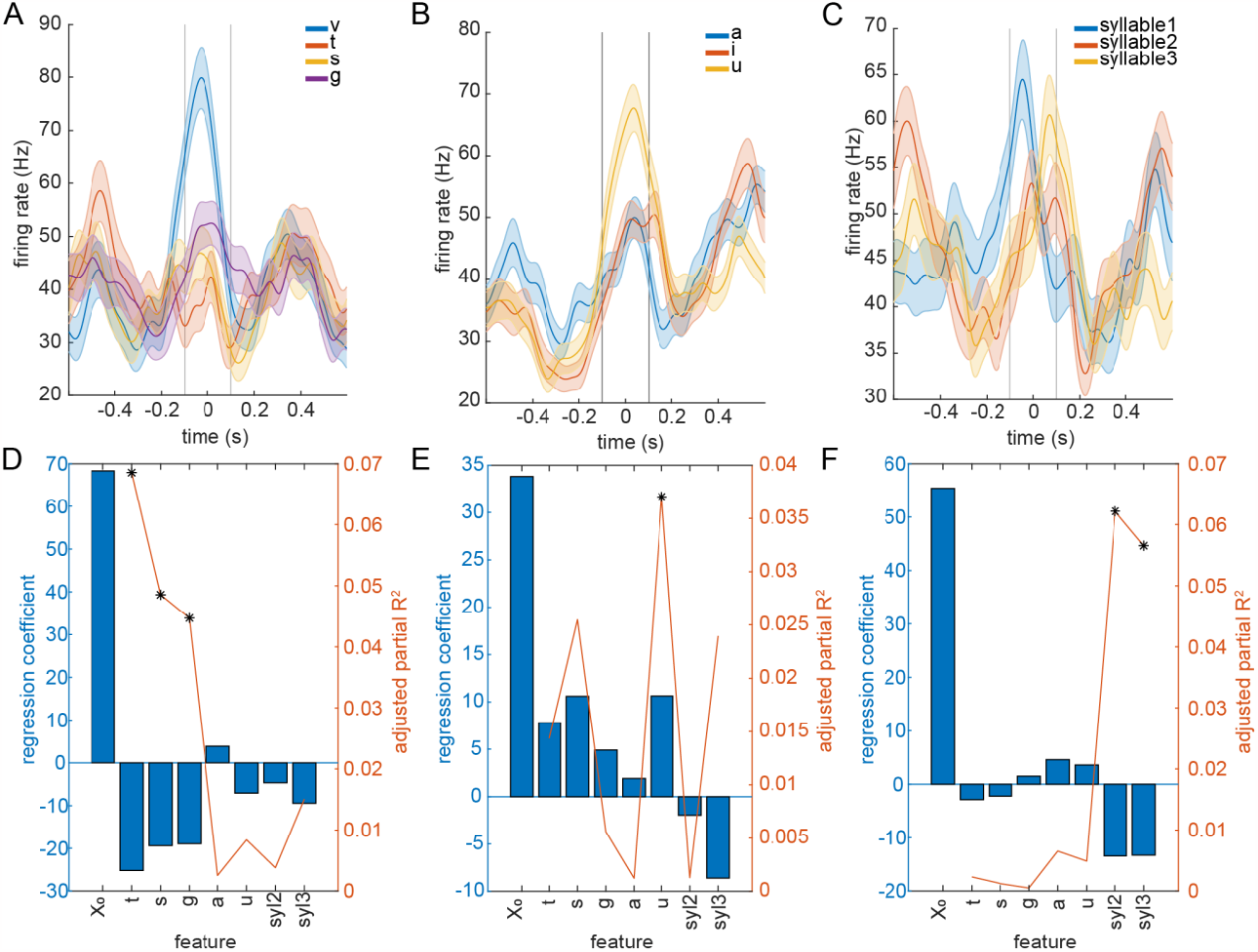
STN neural firing encodes phoneme identity and syllable sequencing. Examples of mean firing rate raster plots indicating consonant (A), vowel (B) and sequence (C) encoding. For each feature category, the spike density function is averaged by feature, across all syllables in a task session. Data are aligned to the consonant-vowel transition (vertical lines depict the 200ms epoch used within which spike activity was fit by the regression model at lag time of 0). (D-F) Regression coefficients (bars) and patrial R2 values (line) for each feature, corresponding to the examples in A-C. Categorical features with a significant contribution to the model (p < 0.05 permutation testing, Bonferroni corrected for 7 features) are marked with an asterisk. X0 corresponds to the reference condition (/vi/ syl1).

**Figure 5.**
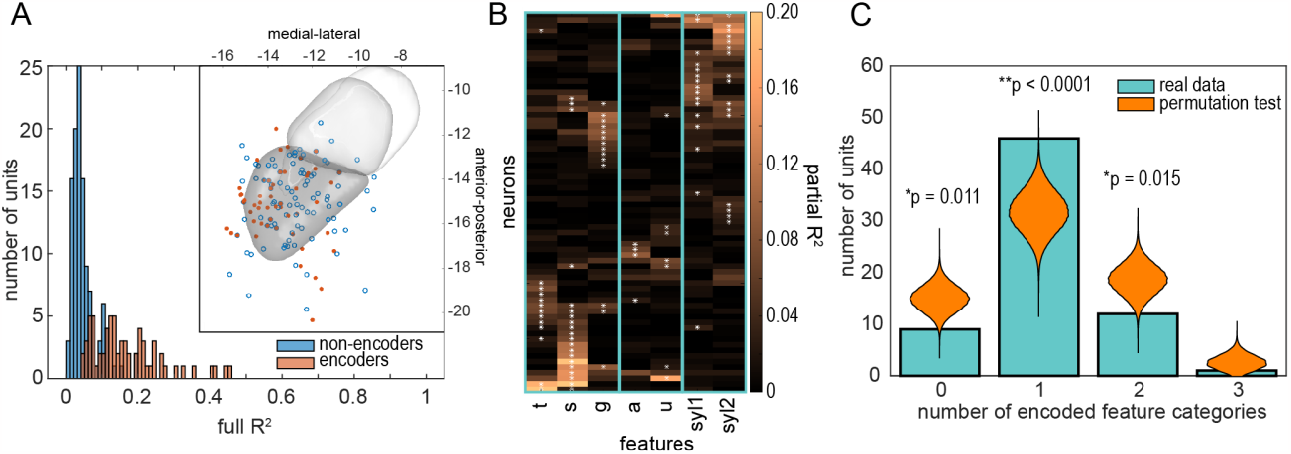
Encoding of phoneme and sequence information by STN neurons is sparse. (A) Distribution of R2 values from the full model fit, for all STN recordings that met the criteria for regression modelling (N = 187). Locations of encoder and non-encoder units (based on permutation testing) are shown in MNI space, along with motor, associative and limbic atlas regions. (B) Partial R2 values corresponding to each categorical variable in the model, for all encoder units, sorted according to a hierarchical clustering procedure. Values corresponding to a significant contribution to the model (p < 0.05 permutation testing, Bonferroni corrected for 7 features) are marked with an asterisk. (C) Units tended to encode speech features exclusively within just one category, with a higher probability then expected by chance (permutation testing), demonstrating sparse encoding of information in this feature space. Significance levels indicate the probability that the real data comes from the distribution produced by the permutation test.

We then identified the specific task factors that contributed to the significant regression effects across the population. Figure 5B summarizes the partial R^2^ values for each task factor (individual vowels, consonants and ordinal positions in the sequence) for all speech-encoding units, with significantly contributing features marked with an asterisk. Visual inspection of Fig. 5B suggests that the distribution of significant partial R^2^ values was markedly inhomogeneous across units. The activity of some units was influenced only by the consonant used (Fig. 5B bottom left), whereas other units were influenced only by the vowel used (Fig. 5B middle) or the ordinal position of the syllable in the 3-syllable sequence (Fig. 5B right top). To investigate this further, we collapsed the partial R2 results for individual features into three general categories of encoding: consonant, vowel and ordinal position. Permutation testing showed that the distribution of encoding across STN units favored exclusive encoding of only one feature category, compared to the distribution expected by chance (Figure 5C).

### Phoneme and sequence feature encoding is independent of modulation type in STN neurons

Neither increase-or decreasetype neurons were more likely to be a speech-encoding unit (41.6% and 39.6% of increase- and decrease-type units respectively) or to encode any of the feature categories examined by the phoneme-sequence model (χ^2^ = 1.13, p = 0.57). Even some neurons whose mean firing rate was not modulated significantly during the triplet repetition task still showed significant encoding of specific features of speech. Speech-related modulation was determined based on an across-trials average of neural activity; therefore, a net zero modulation coupled with a between-trials variability based on speech features can result in significant encoding. However, neurons with no significant modulation in across-trial mean firing rate were less likely to be speech-encoding units (15.9%, 19 of 119 non-modulated units).

### STN neurons encode phoneme features at the time of speech production

To examine the timing of phoneme encoding in the STN, the reduced phoneme model was evaluated based on each unit’s spike density at different lag times relative to the consonant-vowel transition of the produced syllable. We found that 47 of 187 units (25%) that met model criteria encoded phoneme features of speech within the examined epoch (-1.5 to 1.5 seconds relative to each syllable’s consonant-vowel transition). Figure 6A shows the distribution of lag times at which phoneme encoding was observed. Phoneme encoding was most common across neurons (55%, 26 of 47 units, Figure 6A peak in blue histogram) at 25 ms after the consonant-vowel transition. Interestingly, the timing of encoding differed for that of consonants and vowels. The incidence of consonant encoding peaked at -50 ms before the consonant-vowel transition (47% of units) whereas that of vowel encoding peaked at a lag of +175 ms after the consonant-vowel transition (30% of units). Of phoneme-encoding units, 30 encoded only one of the two feature categories (consonant or vowel), whereas 17 encoded features within both categories at some point in the examined epoch. These results observed at the population level were also evident in the task-related modulation of single-units. Figure 6B shows an example of a single-unit that encoded only consonant identity at the range of lag times at which the model was evaluated; Figure 6C illustrates a case in which one single-unit encodes both consonant and vowel identity with vowel encoding reaching a peak at approximately a 300 ms delay after the peak in consonant encoding.

**Figure 6.**
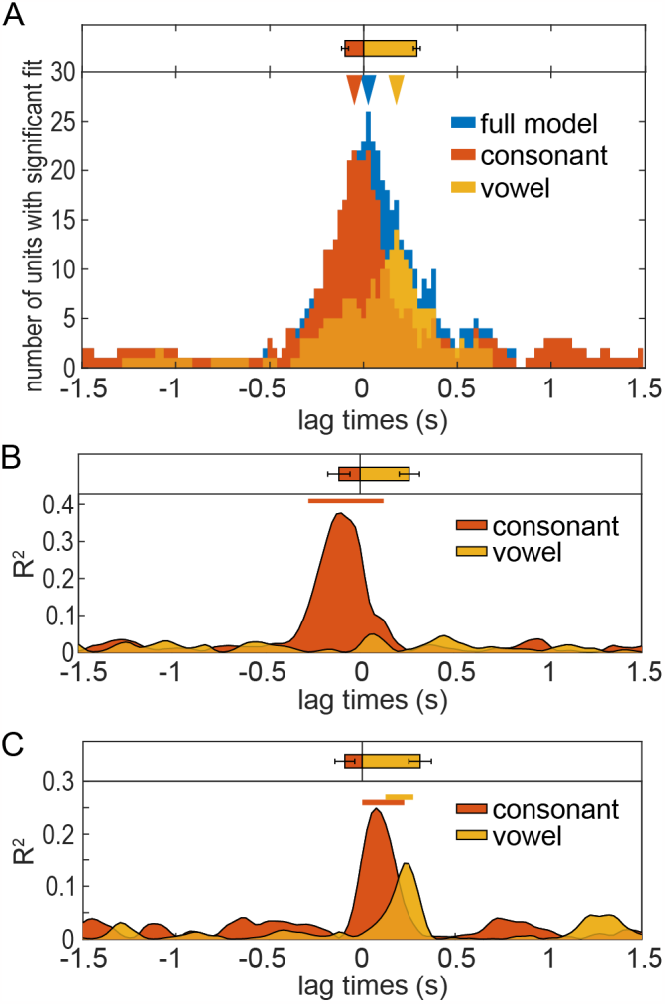
STN neurons encode phoneme features at the time of speech production. (A) Temporal distribution of significant encoding properties for consonant and vowel features. Peak population encoding of each phoneme category occurred at the time of its production. Examples of partial R2 values as a function of lag time for a unit encoding exclusively consonant features (B) and a unit encoding features of both consonant and vowels (C). Histograms showing the mean timing of consonant and vowel production are shown above each plot. Phoneme production timing is averaged across units in (A) and across trials in B-C.

To further characterize the timing of phoneme encoding relative to speech production, we focused our attention on units that encoded consonant identity in the phoneme regression model. We examined firing rate fluctuations of these neurons around the time of acoustic speech onset, which allowed better temporal resolution compared to the modeling approach. While the phoneme model consonant partial R^2^ averaged across these units peaked around the time of acoustic speech onset (Figure 7A), there was considerable variance in the timing of consonant partial R^2^ peaks of individual consonant encoding units. This suggests that the timing of firing rate modulations reflecting consonant encoding is variable across units. Indeed, we found what while some consonant-encoding units showed modulations in activity that began well before the acoustic onset of speech (Figure 7B), others showed changes late, after the acoustic production of the consonant (Figure 7C).

**Figure 7.**
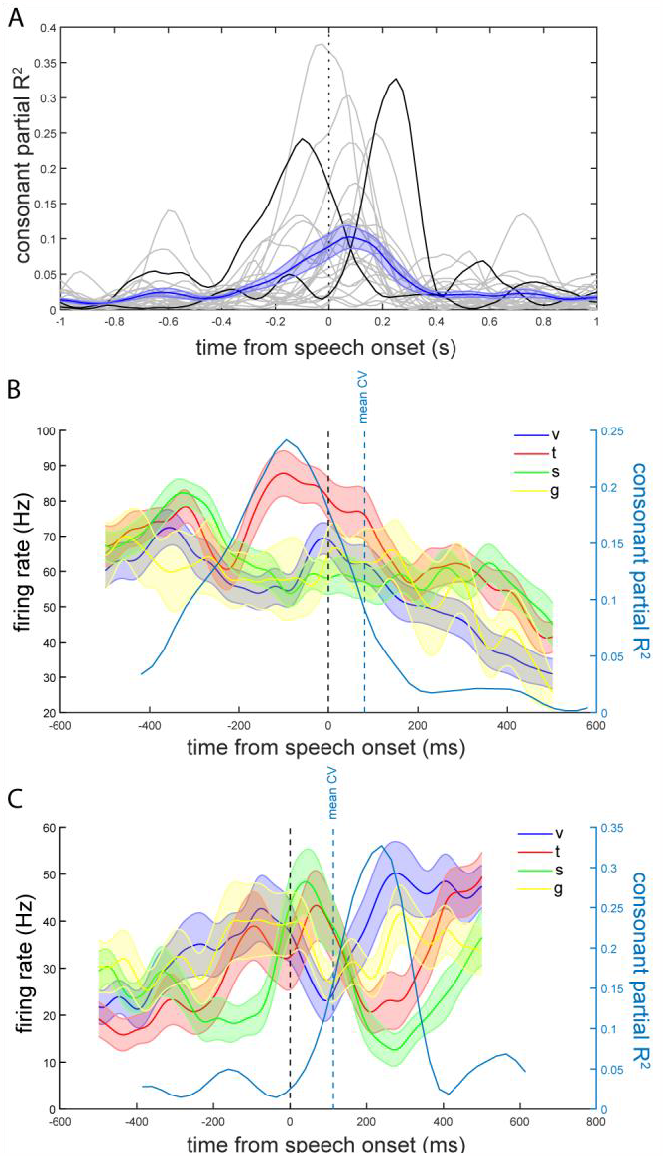
Inconsistent timing of consonant encoding. (A) Population mean and standard error (blue line and shaded region) consonant partial R2 values for regression models evaluated at different lag times relative to the consonant-vowel transition indicate that maximum consonant encoding within the population occurs after acoustic speech onset. Consonant partial R2 values for individual units are overlaid in gray and black; units whose R2 curves are highlighted in black are also shown in panels B and C. Only neurons that showed significant consonant encoding with a minimum of 8 trails containing each consonant in the first syllable position (N=24) were included in this analysis. (B, C) The timing of consonant-related differences in firing rate was inconsistent across units. We observed some units showing consonant encoding prior to the acoustic onset of the first syllable (B), while other units showed encoding well after consonant production (C). Mean firing rate plots (left axis) are aligned to first syllable onset (black dashed line), while consonant partial R2 plots (right axis) are computed based on the regression model lagged relative to the consonant-vowel transition (blue dashed line).

### Spike signal-to-noise is greater than the magnitude of possible acoustic contamination

The mean spike SNR across all units (N=219) considered in this study was 2.07 ± 0.06 (mean ± standard error), while the mean speech-related voltage change SNR across all task sessions was 1.030 ± 0.006, representing a 2-fold difference. This finding supported the visual observation that spike action potentials were not contaminated by artifactual oscillations during speech (see Figure 1B).

## 4. Discussion

In this study, participants repeated aloud syllable triplets played through earphones, a paradigm that allowed investigation of the representation of articulatory and linguistic features in the spiking activity of single-units in the STN. We found that STN neuronal firing encodes specific features of speech, including features related to phoneme identity, as well as information about the ordinal position of the phoneme in a triplet sequence.

Remarkably, a large majority (68%) of STN neurons sampled in this experiment showed a modulation of firing rate during the speech task, including increase (50%) and decrease (30%) type responses. This result is consistent with our previous findings of speech-related changes in STN activity during a reading aloud task, where we observed 53% of recorded neurons with a significant modulation of activity (Lipski et al., 2018). One interpretation of these results is that the population of neurons whose activity is modulated corresponds to a part of an articulator representation within the STN that is engaged by the task. That idea would be consistent with previous neurophysiological studies in humans and animals showing a somatotopic representation of limb movements within the STN (Wichmann et al., 1994; Rodriguez-Oroz et al., 2001; Abosch et al., 2002; Starr et al., 2003; Theodosopoulos et al., 2003; Romanelli et al., 2004; Schrock et al., 2009); it should be noted, however, that no somatotopic pattern of encoding was found in this study. On the other hand, it seems unlikely that such a large proportion of STN neurons would correspond to the representation of speech-related movements. Indeed, in the present study, a smaller proportion of neurons (46%) produced activity that was well fit by the phoneme-sequence model, and only 22.5% of neurons specifically encoded phoneme-related information, which can be described in terms of articulator movements. This finding raises the possibility that many of the modulations in STN activity observed here reflected other processes independent of the control of articulators. Indeed, 14% of recorded neurons encoded the ordinal position of the phoneme in the triplet sequence independent of what specific phoneme was spoken. The idea that some regions of the basal ganglia process information in a modality independent fashion is supported by finding that language syntactic manipulation and tool use share common fMRI activation patterns (Thibault et al., 2021).

Encoding of phoneme and sequence features of speech in the STN was independent of modulation type (i.e. increaseor decreasetype neurons). Based on the firing rate model of the basal ganglia (Albin et al., 1989; DeLong, 1990), increased activity of STN neurons has a suppressive effect on thalamo-cortical targets via the inhibitory GABAergic output neurons of the globus pallidus internus and substatia nigra pars reticulata. The fact that the STN also innervates the globus pallidus externus adds to the complexity of understanding STN function. Given that phoneme and sequence information is encoded via both increases and decreases in STN firing, the STN’s role in speech production appears to involve disinhibition of some motor programs and suppression of others, as is the case for limb movements (Wichmann et al., 1994; Williams et al., 2005).

We found that encoding of speech features was sparsely distributed among STN units such that individual single units tended to encode just one feature category (consonant, vowel or sequence) more often than would be expected if feature encoding was distributed uniformly among units. This finding supports the idea that the transfer of information through the STN specific to individual functions is segregated largely into distinct neural circuits rather than being widely distributed across STN neurons. It is also consistent with previous observations of a segregation of information between subpopulations of STN neurons with respect to both somatotopy (Wichmann et al., 1994; Iwamuro et al., 2017) and cognitive factors (Isoda and Hikosaka, 2008; Pasquereau and Turner, 2017; Nougaret et al., 2022).

We investigated the timing of phoneme encoding by evaluating the phoneme model at different lag times relative to speech production. Encoding of phoneme-related information was most common in STN unit activity coincident with the time of the acoustic production (Figure 6, Figure 7A), which may reflect efference copy-or sensory-related processing (Crapse and Sommer, 2008) rather than advance planning of speech motor output. Sensory functions could include processing of the acoustic signal, which would necessarily take place following the acoustic onset of speech – or somatosensory processing related to articulator positions, which would presumably be ongoing throughout the task. Activity related to efference-copy circuits could be detected as early as the corollary discharge or during later stages of processing. The results of the experiments presented in this study cannot categorically distinguish between these possibilities. Our focused analysis on the timing of consonant-related differences in activity in a subset of units found individual cases in which a single-unit’s consonant encoding appeared well in advance of the onset of speech and others in which that encoding appeared only after the offset of the consonant. We found no consistent pattern in the timing or direction of firing rate changes of these neurons relative to acoustic onset, suggesting that acoustic sensory processing is not the main driver of phoneme-specific activity in the STN. Rather, the phoneme-specific activity described here is likely to reflect STN involvement in a variety of different speech-related processes, which could include advance planning, monitoring of performance during speech and evaluation of performance following phoneme offset, as well as sensory processing.

Experiments described here have been performed with patients with PD, a disorder characterized by abnormal basal ganglia function. It is important to emphasize this caveat when generalizing our findings to the speech function of healthy individuals. However, there are reasons to believe that intracerebral electrophysiology data collected in PD patients will contribute to our general knowledge about how the brain produces speech. There is a substantial history of basic research involving PD patients in which performance in the disease state has been used to inform models of normal function. Published data aligns well with related findings and expectations from the literature (Kondylis et al., 2016; Lipski et al., 2017, 2018), suggesting that this approach can reveal general principles that are not specific to the disease state. It is important to note that DBS surgery offers the only opportunity to record from the STN while a person is speaking, as currently there is no other routine clinical indication for implantation of electrodes into basal ganglia nuclei.

Studies from our group (Bush et al., 2022) as well as others’ (Roussel et al., 2020; Berger et al., 2022) show that intracranial electrophysiological recordings are sensitive to contamination from acoustic vibrations during speech. However, we have found that in our hands, such contamination of microelectrode signals does not interfere with the sorting and isolation of large-amplitude extracellular spikes used in this study, because properly sorted extracellular spikes have features that are distinct from an audio artifact, including: (1) a larger amplitude of the spike waveform; (2) a stereotyped waveform shape that produces distinct clusters in principal component space during the sorting procedure; and (3) presence of a refractory period visible in the inter-spike interval histogram in single units. We show that the signal-to-noise ratio of spikes in these experiments exceeds that of possible acoustic contamination by a factor of two.

In summary, our findings implicate the STN in two separate speech-related processes – one related to the sequencing of speech vowel-consonant pairs, independent of articulatory features, and the other related to the phonemes being produced. Those two relationships substantiate the involvement of basal ganglia circuits at multiple levels of the neural control of speech. These findings advance our understanding of neural substrates of human speech processing and shed light on potential involvement of the STN in other complex human behaviors.

